# Testosterone Changes in Female Pandas in Estrus

**DOI:** 10.1101/863704

**Authors:** Xuefeng Liu, Lei Yuan, Wei Wang, Yuyan You, Yanhui Liu, Tao Ma, Zezhong Wang, Xuelin Jin, Qingyi Ma, Yanping Lu, Jinguo Zhang, Sufen Zhao

## Abstract

Currently, the majority of giant panda breeding is carried out by cage-mating or artificial insemination based on estrogen levels and behavior in female pandas. However, studies have shown that testosterone levels both in women and in non-human primate females have a significant effect on the desire to mate. In this study, we wanted to explain how testosterone levels of female giant pandas would change during estrous. In this study, 23 accounts of rutting were recorded in 10 female pandas from 2009 to 2012. Changes in urinal testosterone levels were monitored and compared with estradiol values. Our data showed that, for female pandas in estrus, testosterone levels after the estradiol peak was significantly higher than before, and the testosterone peak occurred 4 days after the estradiol peak. Furthermore, testosterone and estradiol level were only significantly correlated after peak estradiol levels peaked, and not before. Finally, out findings suggest that testosterone could help us better understand hormone variation during panda estrus, as well as help aid in the natural breeding of pandas.

## 1. Introduction

Giant pandas (*Ailuropoda melanoleuca*) are rare in the world. Under artificial breeding conditions, a healthy female giant panda experiences one estrus period per year, from February to June. The duration is short and rutting behavior is usually observed. As estrus approaches, the vaginal orifice becomes red and swollen and the female may rub the genital region with a paw or on objects (Anon, 1974; Kleiman et al., 1979).

In recent years, many researchers studied changes in giant panda reproductive endocrine (Zeng et al., 1984; Shi et al., 1988; Peng et al., 1993; Xie et al., 1993) as well as variation in related hormones such as estradiol, progesterone, lutinizing hormone, etc. (Bonney et al., 1982; Hodges et al., 1984; Liu, 1988; Zeng et al., 1990; Li et al., 1993; Yu et al., 2003). These studies closely examined the relationship between hormones and ovulation. However, other studies found that testosterone levels in women and in non-human primate females also have a significant effect on the desire to mate (Everitt et al., 1971; Davis and Tran, 2001; Gao et al., 2007). Previous research has also shown that testosterone levels in women and sexual readiness showed a significant positive correlation (Sherwin et al., 1987; Van et al., 1997; Gumell and Chatterjee, 2001). According to a study of Sichuan golden monkeys during their breeding period, solicitous behavior in three females showed significant positive correlation with their testosterone levels (Gao et al., 2007).

In our study, a radioactive immunity method was to monitor changes in testosterone during female giant panda breeding, and is compared with changes in estradiol. This research provides reference data for better understanding of the hormonal changes during the panda’s estrus breeding period.

## 2 Materials and Methods

### 2.1 Ethics statement

The study was approved by the Beijing Municipal Committee of Animal Management before sample collection.

All experiments were performed in accordance with the approved guidelines and regulations.

### 2.2 Materials

Experimental animals included 10 female pandas from the rescue breeding research center in Shanxi and Beijing Zoo. From 2009 to 2012 urine was collected and examined a total of 23 times (Table 1). Urine was only measured after estrus behavior was observed. Estrus behavior in giant female pandas was defined by Bonney et al (1982). During the female estrus period, urine was collected between 8:00 to 10:00 A.M. every day and immediately stored in a −20°C freezer. Day 0 is defined as the day of peak estradiol. One urine sample was collected daily from each animal starting 8 days before and 8 days after Day 0 for a total of 16 days.

**Table 1.** Information of sampling.

| Studbook | Name | Birth year | Sampling year | Institution |
| --- | --- | --- | --- | --- |
| 320 | Lele | 1986 | 2009, 2011 | Beijing Zoo |
| 403 | Jini | 1993 | 2010, 2011, 2012 | Beijing Zoo |
| 566 | Yinghua | 2003 | 2010, 2011, 2012 | Beijing Zoo |
| 652 | Mengmeng | 2006 | 2010, 2011 | Beijing Zoo |
| 444 | Xuexue | 1988 | 2009 | The Rescue Breeding Research Center in Shanxi |
| 509 | Zhuzhu | 2000 | 2009, 2011, 2012 | The Rescue Breeding Research Center in Shanxi |
| 562 | Yangyang | 2003 | 2009, 2011, 2012 | The Rescue Breeding Research Center in Shanxi |
| 603 | Xinxin | 2005 | 2011, 2012 | The Rescue Breeding Research Center in Shanxi |
| 660 | Niuniu | 1997 | 2009, 2011, 2012 | The Rescue Breeding Research Center in Shanxi |
| 699 | Chengcheng | 2006 | 2012 | The Rescue Breeding Research Center in Shanxi |

### 2.3 Urine sample processing

Fresh urine samples are tested immediately. Cryopreserved urine was placed in ice baths and tested after it reached room temperature.

### 2.4 Hormone detection

Using the radioactiveimmunity method described by Monfort et al. (1989), with some slight modifications. The radiation immunoassay reagent kit used was from Beijing Kemeidongya Biological Technology Co. Ltd. and the GC2010 Ria Gamma Counting Instrument from Hefei ZhongCheng Co. were used for testosterone and estradiol measurement. Urine creatinine detection was performed with the creatinine (picric acid method) kits from Beijing Kemeidongya Biological Technology Co. Ltd. and the 7080 biochemical analyzer from Japan Hitachi.

### 2.5 Data processing

Testosterone and estradiol measurement data were used for creatinine values for calibration. In order to eliminate possible errors, hormone concentrations with ria determination results and the ratio of creatinine content of the same sample. Statistical procedures were performed using SPSS 19.0.

## 3 Results

Statistical analysis was performed using the results of the 23 cases of rutting where the female giant pandas’ testosterone and estradiol level (Mean ± SE) and production curve were calculated (Fig. 1). Figure 1 shows that during the giant panda’s estrus breeding period, in urine, the testosterone levels peaked 4 days after estradiol levels peaked. For the spearman rank correlation test, testosterone and estradiol level were not significantly correlated before peak estradiol, while after that they were significantly correlated (Before: *n* = 118, *r* = 0.124, *P* = 0.182; After: *n* = 99, *r* = 0.239, *P* = 0.017). Furthermore, testosterone levels were significantly higher after estradiol peaked than before (Independent-Samples t Test: *n* = 250, *t* = 1.975, *P* = 0.044).

**Fig. 1.**
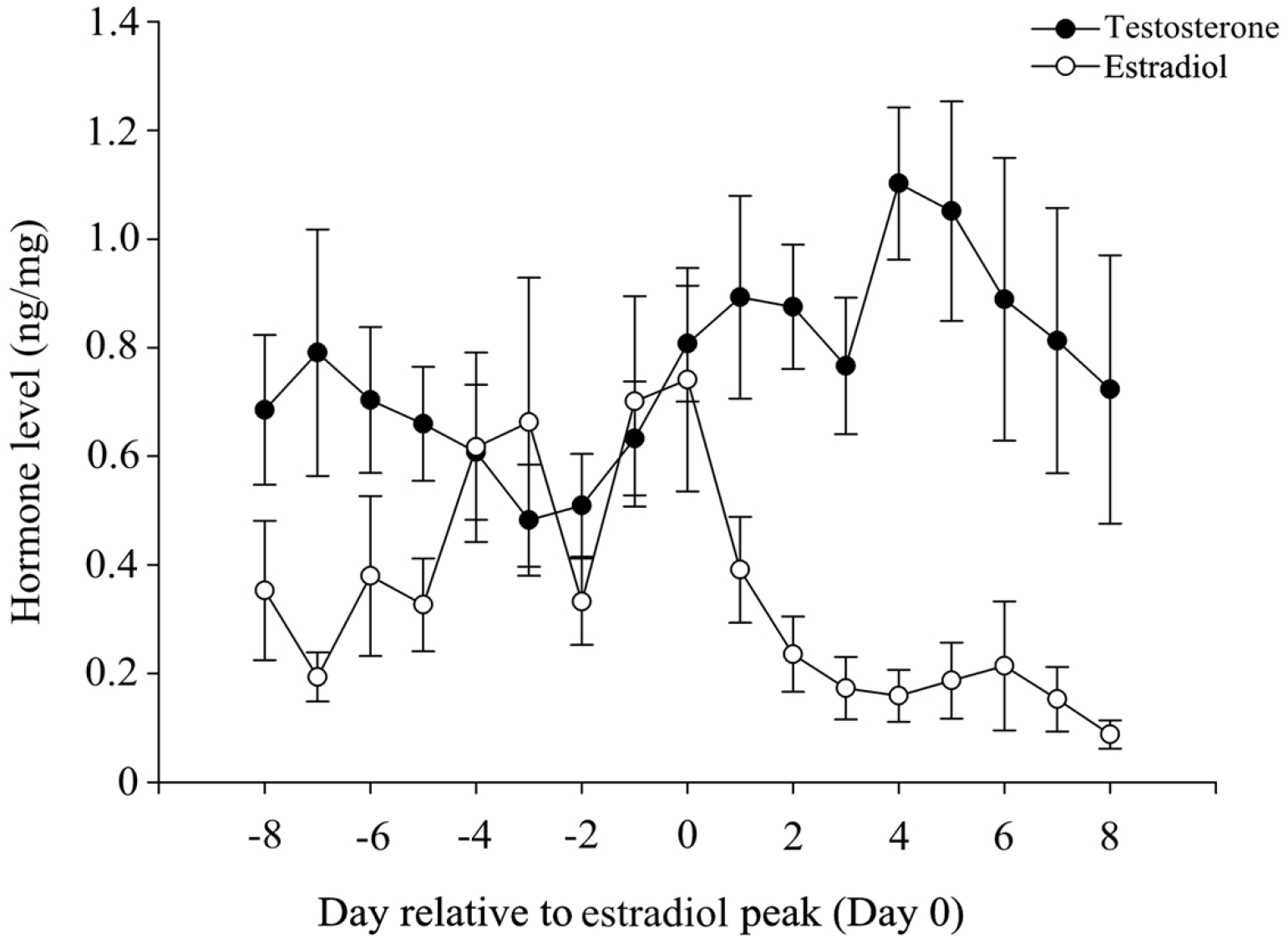
Levels of Urinary Testosterone and Estradiol of Female Giant Pandas during Estrus

## 4 Discussion and Conclusions

This was the first study to monitor variations in testosterone levels in female giant panda during estrus. We observed that its changes were similar to that of estradiol. Previous research in animal reproductive endocrinology showed that testosterone is a precursor of estradiol and testosterone can be converted into estrogen by aromatase. For pandas in estrus, urinary estradiol peaks earlier than testosterone by 4 days (Fig. 1). This may be due to the sudden increase of ovarian secretion of testosterone or to the aromatase conversion function reduction. Whether or not this phenomenon is associated with ovulation is worthy of further research.

Some research indicate that optimal breeding time should be on the following day or even on the third day after the peak (Hodges et al., 1984; Zeng et al., 1984). Natural breeding of pandas also occurs after estradiol peaks (Peng et al., 1993). Furthermore, research has also shown that testosterone levels in females and sexual readiness showed a significant positive correlation (Sherwin et al., 1987; Van et al., 1997; Gumell and Chatterjee, 2001). According to a study of Sichuan golden monkeys during their breeding period, solicitous behavior in three females showed significant positive correlation with their testosterone levels (Gao et al., 2007). In this study, we found that testosterone levels are significantly higher after estradiol levels peak. Therefore, testosterone may be a new indicator to evaluate the natural breeding of pandas and provide a reference for further understanding hormone variation during panda fertility.

## Acknowledgement

This study was funded by the U.S. Memphis Zoo’s Protection Project for Giant Pandas. Thanks to the Beijing Zoo and its staff in the Protection of the Rare Wildlife Rescue Breeding Research Center in Shanxi Province for providing research support.

